# Recycling of energy dissipated as heat accounts for high activity of Photosystem II

**DOI:** 10.1101/842591

**Authors:** Monika Zubik, Rafal Luchowski, Dariusz Kluczyk, Wojciech Grudzinski, Magdalena Maksim, Artur Nosalewicz, Wieslaw I. Gruszecki

**Affiliations:** Department of Biophysics, Institute of Physics, Maria Curie-Sklodowska University, 20-031 Lublin, Poland; Institute of Agrophysics, Polish Academy of Sciences, Doswiadczalna 4, 20-290 Lublin, Poland

## Abstract

Photosystem II (PSII) converts light into chemical energy powering almost entire life on Earth. The primary photovoltaic reaction in the PSII reaction centre requires energy corresponding to 680 nm that is significantly higher than in the case of the low-energy states in the antenna complexes involved in the harvesting of excitations driving PSII. Here we show that despite seemingly insufficient energy, the low-energy excited states can power PSII thanks to the activity of the thermally-driven up-conversion. We demonstrate the operation of this mechanism both in intact leaves and in isolated pigment-protein complex LHCII. A mechanism is proposed, according to which the effective utilization of thermal energy in the photosynthetic apparatus is possible owing to the formation of LHCII supramolecular structures, leading to the coupled energy levels, corresponding to approx. 680 nm and 700 nm, capable of exchanging excitation energy through the spontaneous relaxation and the thermal up-conversion.

**TOC GRAPHICS:** 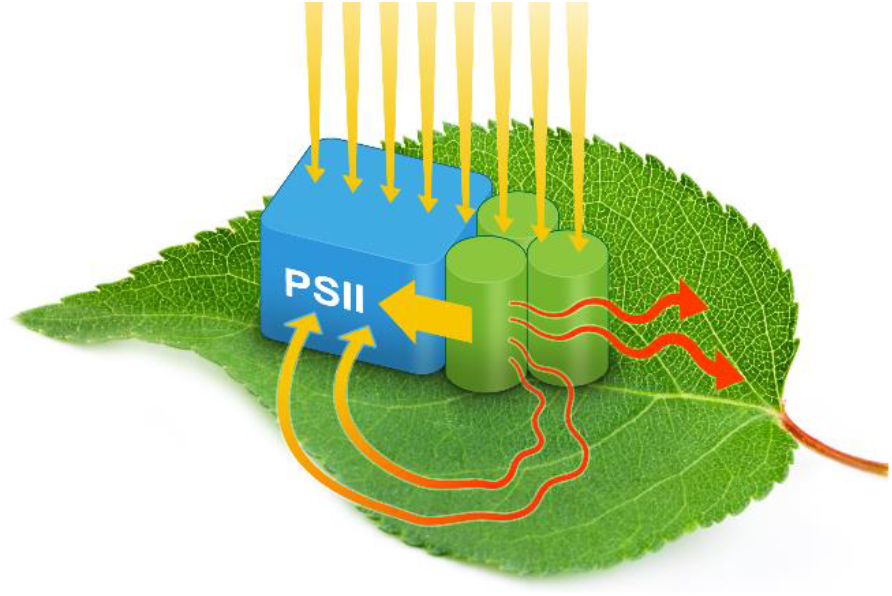

---

Life on Earth is powered by sunlight and photosynthesis is practically a sole process able to convert the energy of electromagnetic radiation to the forms which can be directly utilized to drive biochemical reactions in living organisms ^1^. Importantly for life in our biosphere, oxygenic photosynthesis supplies molecular oxygen to the atmosphere, which most of the organisms use for respiration ^2^. Photosynthetic oxygen evolution is directly associated with the activity of Photosystem II (PSII), in which electrons can be detached from water molecules ^3^ due to the relatively high redox potential of the P680^•+^ radical cation created by a photoreaction in the reaction center ^4^, estimated to be 1.26 V ^5^. PSII has originally evolved 2.5 billion years ago and is located in the photosynthetic membranes of cyanobacteria, algae and plants ^6^. Oxidation of water would not be possible in Photosystem I (PSI) –type complexes present in all photosynthesizing organisms due to the insufficient redox potential of the P700^•+^ radical cation created by a photoreaction in its reaction center. Most of the photosynthetic pigments in plants, energetically coupled to PSII and PSI contribute to the separate chlorophyll *a* (Chl *a*) fluorescence emission bands centering respectively at ca. 680 nm and 735 nm ^7, 8^ (see Fig. 1a). Intriguingly, at the low temperature, the fluorescence emission from Chl *a* molecules associated with PSI is substantially more intensive than from the pool associated with PSII. The opposite proportion can be observed at room temperature, in accordance with the numerous reports ^8^. As it might be expected, the decrease in the temperature, from the room temperature to 77 K, resulted in the increase in Chl *a* fluorescence yield both in PSII core complexes and in PSI ^8^. Importantly, the factor of increase in fluorescence quantum yield was substantially different in the case of PSII and PSI, namely, about 2 versus 20 respectively ^8^. A relatively low enhancement at 77 K of a fluorescence signal from the Chl *a* spectral forms energetically coupled to PSII, as compared to PSI is not understood and may be considered peculiar from the standpoint of photo-physics. Basically, there are several possible interpretations of this observation. The most direct interpretation could be that this is a manifestation of the impaired energetic activity of PSII at low temperatures ^8^. While this interpretation is far from speculating, it does not provide any explanation based on the specific molecular mechanisms responsible for the observed differences. Regarding possible molecular mechanisms, one may consider a different proportion between the radiative and non-radiative energy dissipation channels of excited Chl *a* energetically coupled to PSII and PSI, at high and low temperatures. This mechanism, though likely, still appears to be proposed at a relatively high level of generality. Experiments presented below, designed to investigate such a possibility, led us to unveil the process of thermally-driven up-conversion operating in the photosynthetic systems, as one of the highly probable mechanisms responsible for this observation. In the present study, the integration of the Chl *a* fluorescence emission spectra in leaves in the spectral region representing mostly PSII (wavelengths below 703 nm) and in the spectral region representing largely PSI (wavelengths higher than 703 nm) gives the ratio of photons emitted by PSII and PSI as high as 1.16 ± 0.22 at 298 K but only 0.20 ± 0.02 at 77 K (mean value from three different leaves ± S.D., Fig. 1a). Such a result goes along with the general observation regarding the relatively low photophysical activity of PSII at low temperatures ^8^. Upon the temperature decrease, the low-energy spectral forms are enhanced to a higher degree than the high-energy spectral forms. This suggests that photophysical mechanisms such as thermally-driven up-conversion can be potentially involved in shaping the emission spectrum at room temperature. Such a situation corresponds directly to the Chl *a* fluorescence emission spectrum recorded from leaves, at the low and high temperatures (Fig. 1a). It is therefore very likely that thermal activation of the low-energy states, corresponding to wavelengths higher than 700 nm, is necessary for effective excitation of the high-energy spectral forms supplying energy to the P680 reaction center of PSII. Such a process cannot take place at 77 K due to insufficient thermal energy. In principle, a process of the thermally-driven up-conversion can operate effectively in the photosynthetic apparatus of plants owing to the fact that the overall energy conversion efficiency in photosynthesis does not exceed 6 % and most of the energy of absorbed light quanta is dissipated as heat ^9, 10^. According to the estimation by Jennings et al. heat accounts for 95-98 % of absorbed light energy released into the environment ^9^. The major fraction of heat released to the environment is associated with relaxation of the singlet-excited electronic states of chlorophylls corresponding to the Soret spectral region to the Q_y_ level representing the red spectral region. A direct demonstration of the operation of the heat-driven uphill energy transfer in the photosynthetic apparatus in leaves is shown in Fig. 2.

**Figure 1.**
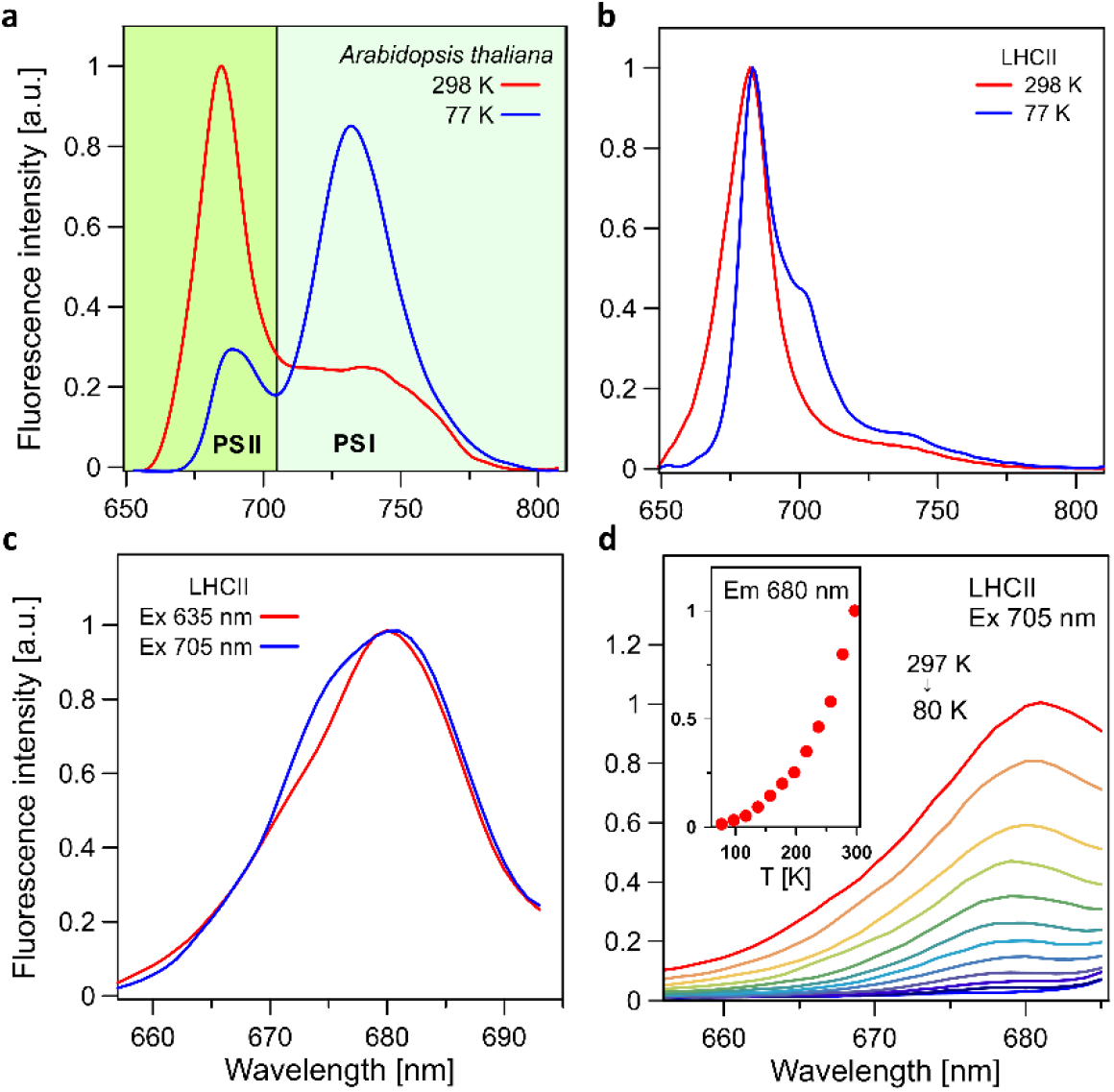
(a) Fluorescence emission spectra recorded from *A. thaliana* leaves. Excitation at 635 nm. The spectra were recorded at 77 K and 298 K, indicated. The spectra were area-normalized and presented on an arbitrary scale. The short-wavelength and the long-wavelength emission bands are assigned to PSII and PSI respectively. (b) Fluorescence emission spectra recorded from the sample containing supramolecular structures of LHCII formed spontaneously in the environment of the lipid membrane. Excitation at 635 nm. The spectra normalized at the maximum. (c) Fluorescence emission spectra recorded from the LHCII sample as in panel b, at 298 K with the laser excitations set at 635 nm or 705 nm, indicated. (d) Fluorescence emission spectra recorded from the LHCII sample as in panel b, recorded at different temperatures in the range from 297 K to 80 K. Excitation was set in the lower energy spectral region, at 705 nm. The inset shows the temperature dependence of fluorescence intensity at 680 nm, based on the spectra presented. The set of the spectra is presented on an arbitrary scale.

**Figure 2.**
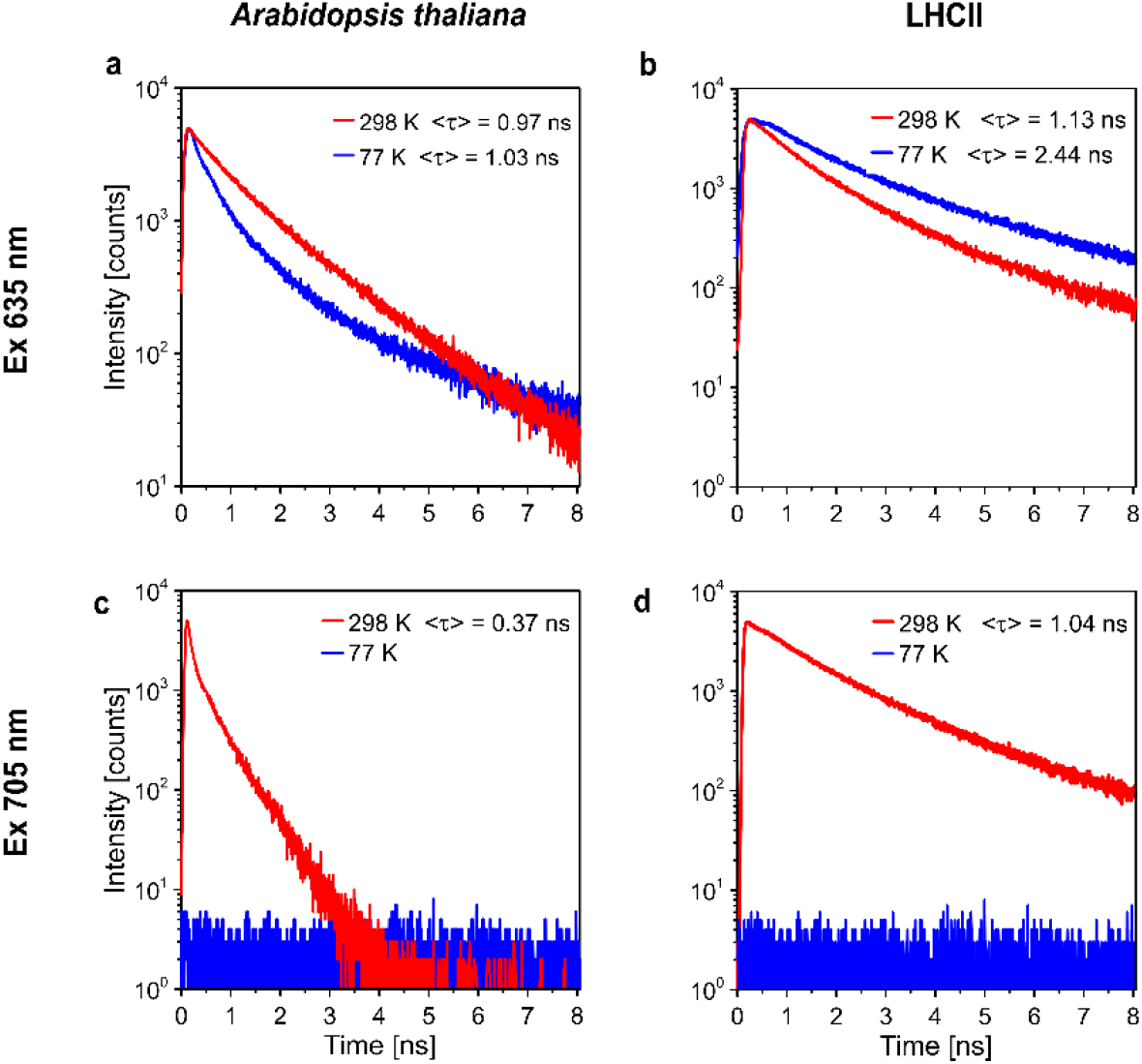
Chlorophyll *a* fluorescence decay kinetics recorded from intact leaves and LHCII samples. The kinetics were recorded from intact leaves of *A. thaliana* (a and c) and from the samples containing supramolecular structures of LHCII formed spontaneously in the environment of lipid membrane (b and d), at 77 K and 298 K, indicated. Two combinations of the excitation and fluorescence observation wavelengths were applied: Ex 635 nm/Em 680 nm (panels a and b) and Ex 705/Em 680 nm (panels c and d). Displayed are amplitude weighted average fluorescence lifetimes (<τ>) calculated for each system. Differences in the Chl *a* fluorescence decay kinetics recorded from intact leaves and LHCII express differences in the complexity of both systems. In particular, fluorescence kinetics in intact leaves at the room temperature is modulated by the activity of the photosynthetic reactions.

Fluorescence of Chl *a* excited in the lower energy region (at 705 nm) and detected at higher energies (at 680 nm) can be detected in intact leaves at the room temperature (298 K) but not at 77 K (Fig. 2c). Efficient and fluent operation of photosynthesis is assured by the activity of pigment-protein complexes, called the antenna, collecting photons and transferring excitation energy towards the reaction centres responsible for the primary electric charge separation ^11, 12^. The largest light-harvesting pigment-protein complex of plants, referred to as LHCII, is particularly well suited to play a photosynthetic antenna function, owing to the relatively high concentration of chlorophylls, the presence of xanthophylls (effectively protecting the complex against photo-damage) and internal pathways of extremely efficient excitation energy transfer within the network of the protein-embedded chromophores ^11, 12^. A consequence of exceptionally high protein crowding in the thylakoid membranes is the clustering of antenna complexes, potentially resulting in excitation quenching and thermal energy dissipation ^13–18^. Molecular organization of LHCII can be modelled in the experimental system composed of isolated LHCII embedded into the lipid membranes formed with chloroplast lipids ^16, 19–21^. All the photosynthetically active chlorophylls are embedded in the functional pigment-protein complexes and, in principle, there are no major differences in the chemical environment in PSI and PSII. On the other hand, spectral properties of the pigments are critically dependent on their molecular organization, including the molecular organization of pigment-protein complexes. LHCII is a major antenna complex in the photosynthetic apparatus of plants, comprising roughly half of chlorophyll molecules in the biosphere. In the present study, we applied a system comprising supramolecular structures of LHCII in the environment of membranes formed with chloroplast lipids, to model the natural environment of the photosynthetic apparatus. A spontaneous self-association of the antenna complexes in such a system gives rise to the low-energy band in a low-temperature Chl *a* fluorescence emission spectrum, centered in the region of 700 nm, accompanying the principal band centered in the region of 680 nm (Fig. 1b), in accordance with the previous reports ^17, 22–24^. The long-wavelength band in this particular spectral region (~700 nm) has been assigned to LHCII clusters in the natural thylakoid membranes ^25^. There are several lines of evidence for the formation of such supramolecular structures of LHCII in the thylakoid membranes of plant chloroplasts, including the one based on the circular dichroism analyses ^26^, low-temperature fluorescence spectroscopy ^25^ and direct imaging based on electron microscopy ^14^. The long-wavelength band centred close to 700 nm can be also resolved at 77 K in isolated thylakoid membranes ^27^ and leaves ^8^. The “700 nm band” is a hallmark of aggregated LHCII ^22–24^ although it can be also detected from the trimeric LHCII particles in single-molecule experiments ^28^. Importantly, the “700 nm band” (referred to as E700) can be resolved exclusively in the fluorescence emission spectra of aggregated LHCII recorded at low temperatures but not at physiological temperatures ^17, 22–25^. This observation suggests possible depopulation of the E700 state by an uphill energy transfer, e.g. to the E680 energy level. Direct evidence for the operation of such a process is presented in Fig. 1c showing a comparison of the fluorescence emission spectra recorded from the same LHCII sample excited in the higher and the lower energy regions (at 635 nm and 705 nm) concerning the emission spectral window. Almost identical shapes of both the emission spectra recorded are consistent with the interpretation according to which both the fluorescence emissions originate from the same energy level (Fig. 1c, see also Supporting Information Fig. S1). Comparison of the quantum yields of fluorescence excited at 635 nm and 705 nm shows that the quantum yield of the emission excited at 705 nm is lower by a factor of 6.3 than the fluorescence excited in the higher energy region (at 635 nm). In principle, the ratio of the uphill and downhill rate constants multiplied by the state populations should follow the Boltzmann distribution ^29^. The difference between the E680 and E700 states (420 cm^−1^) expressed in the kT units corresponds to the temperature of 627 K that is higher than the room temperature (298 K). On the other hand, the energy gap and kT ratio (ΔE/kT) that is only as high as 2.1 at 298 K, makes the uphill energy transfer relatively efficient (exp(-ΔE/kT)=0.12), as manifested by the fluorescence quantum yield of the anti-Stokes excitation. The effective energy gap between the E680 and E700 states is likely to be even lower than 420 cm^−1^ owing to the degeneracy of states at room temperature. The fact that the fluorescence spectrum can be recorded under the anti-Stokes conditions and that the quantum yield is relatively high is a direct manifestation of the activity of the up-conversion process. Fluorescence intensity in such a system, recorded at 680 nm and excited at lower energies, drops down with the temperature decrease (see Fig. 1d) and with increasing a distance between the excitation and observation wavelengths (see Supporting Information Fig. S2). Importantly, application of two-dimensional electronic spectroscopy, which correlates the fluorescence excitation and observation wavelengths, enabled to detect a temperature-dependent uphill excitation energy transfer even in the trimeric LHCII ^30^. Another kind of evidence for the operation of the thermally-driven up-conversion in the supramolecular structures of the complex is shown in Fig. 2d presenting fluorescence decay kinetics, excited at 705 nm and detected in the higher energy region, at 680 nm. The process of a thermally-driven up-conversion can be observed at room temperature but it is not possible to operate at 77 K, for the energy reasons (Fig. 2d). This means that the observed thermally-driven up-conversion has to be combined with a thermal deactivation of chromophores electronically-excited and localized in the close neighborhood. It should be emphasized that the illuminated LHCII proved to be a very efficient emitter of heat that can be transmitted over long distances in the supramolecular structures of the protein ^31^. The fact that fluorescence emission from the E700 can be detected at low temperatures, enables to determine fluorescence lifetime of this state (Fig. 3, Fig. S3). It appears that the average lifetime of the E700 is substantially longer than that of the E680 state: 4.18 ns versus 2.44 ns, very close to the previous determinations of similar LHCII systems ^24^. This suggests that the presence of the E700 energy level, below the E680 state, creates conditions for effective excitation quenching from this latter state, by a downhill energy transfer. Most probably, due to such a quenching, the fluorescence lifetime of E680 is even shorter at the higher temperatures (1.13 ns, see Fig. 2b). Intriguingly, the fluorescence emission from the E700 state of aggregated LHCII is not effectively observed at room temperatures (see Fig. 1b and the literature references ^17, 22–25^). This observation implies that under such conditions the E700 is almost entirely depopulated via non-radiative processes including the uphill energy transfer to E680 and thermal dissipation. It is also possible that the E700 state is depopulated via energy transfer to the low-energy states of LHCII (730-780 nm, see Fig. 1b, Fig. S4). Such a long-wavelength fluorescence emission band is particularly intense at low temperatures, under the absence of the thermally-driven up-conversion. The process of thermally-driven up-conversion demonstrated to operate in the supramolecular structures of LHCII formed spontaneously in the environment of chloroplast lipids, operates also efficiently in the aggregated structures of LHCII formed in a water medium (see Fig. S5, Fig. S6 and Supporting Information Table S1). Interestingly, the long-wavelength spectral forms can be detected in a single molecule emission spectra of LHCII ^28^. Coexistence of the two-direction energy transfer pathways between the E680 and E700 states that are present in supramolecular structures of LHCII, namely the spontaneous down-conversion from E680 to E700 and the thermally-driven up-conversion from E700 to E680, can be discussed in the context of overall excitation energy flows in the photosynthetic apparatus of plants (see the model presented in Fig. 4).

**Figure 3.**
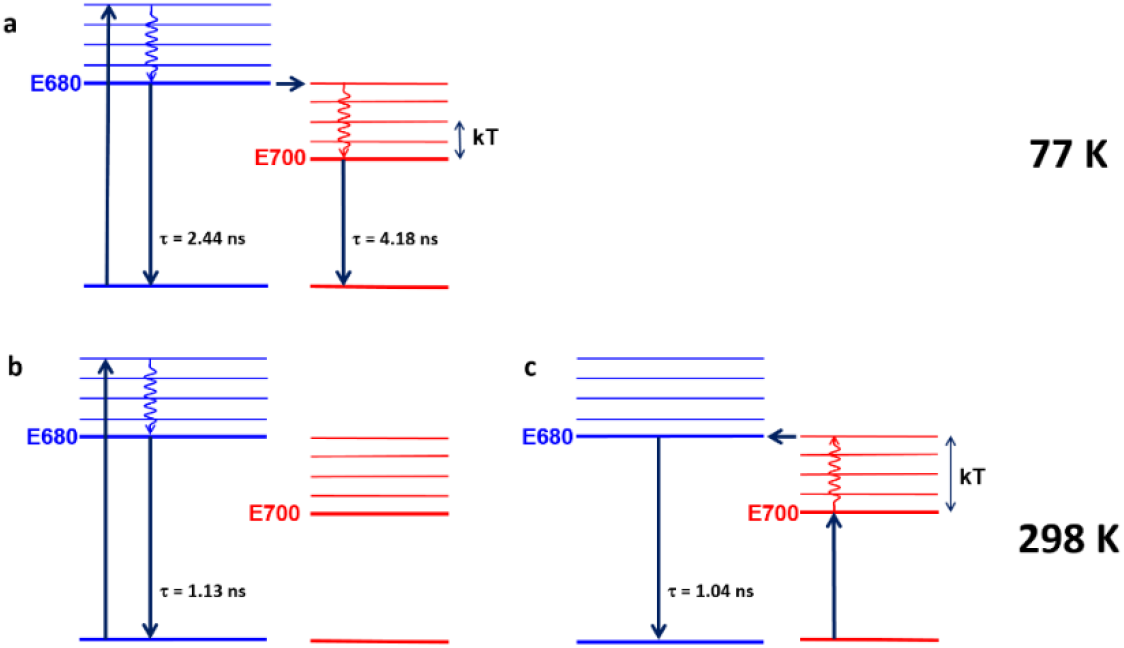
Energy level diagrams representing selected electronic states and transitions in supramolecular structures of LHCII. The average fluorescence lifetime values are also reported, determined based on the decay kinetics shown in Fig. 2 and Fig. S7. Vertical arrows up represent light absorption, vertical arrows down represent fluorescence, wavy and horizontal arrows represent nonradiative processes. The examples of a multiexponential analysis of Chl *a* fluorescence decays in LHCII are presented in Fig. S7.

**Figure 4.**
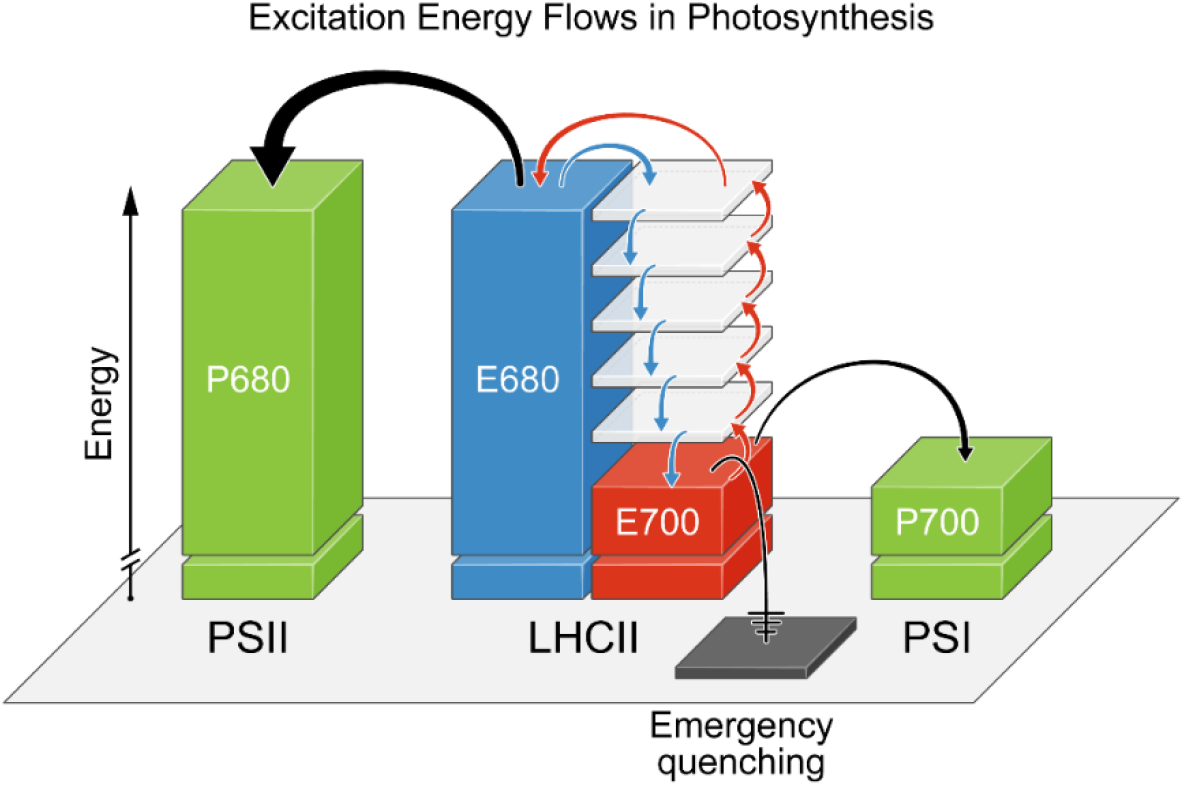
A simplified “energy recycling unit” model representing the excitation energy flows in the photosynthetic apparatus of plants. The scheme presents the excitation energy flows between the light-harvesting antenna complexes (LHCII) and the reaction centres of Photosystem II (P680) and Photosystem I (P700). Two experimentally identified, energetically coupled excitation energy states of supramolecular structures formed by LHCII are marked as E680 and E700. Energy circulation between these states is represented by arrows: the quenching of E680 by E700 and the thermally-driven up-conversion from E700 to E680. An emergency excitation quenching channel leading to thermal dissipation of excess excitation energy is also shown.

The central element of this model, which may be referred to as the “energy recycling unit”, is based upon spontaneously formed supramolecular structures of LHCII. We have modelled the formation of such structures in LHCII-containing lipid membranes. Dependently of an actual protein-lipid ratio, the proportion between the E680 and E700 bands can be different in the fluorescence emission spectra, reflecting ability of LHCII to self-aggregate in a system ^17^. One of the weaknesses of the model applied is a possibility of incorporation of the protein in two opposite orientations, i.e. the N- and C-termini facing to both sides of the lipid bilayers, in contrast to the native thylakoid membranes. On the other hand, very close agreement of the spectral shapes representing the aggregated structures of LHCII formed in the natural thylakoid membranes and under laboratory conditions justifies the application of such a system for model study ^25^. Nevertheless, for the sake of caution, the conclusions regarding the thermal up-conversion, drawn from experiments performed on model membranes, were confirmed in this study by the results of the experiments carried out on intact leaves.

Numerous consequences of the operation of the molecular system based on supramolecular structures of LHCII can be envisaged, important for the activity of PSII and the efficiency of photosynthetic energy conversion in plants. The main consequence seems to be the ability to use low-energy excitations (corresponding to wavelengths higher than 680 nm) to drive the photo-physical processes in the PSII reaction centre, thanks to the utilization of a fraction of the excitation energy dissipated as heat in the pigment-protein antenna systems. Concerning the utilization of near-infrared radiation, it is worth mentioning that the core complex of PSII also contains some red-shifted states, in the core antenna CP43/CP47 and possibly some states emitting light up to more than 700 nm, which might be part of the reaction center excimer and less fluorescent. The 695 nm band was identified in the low-temperature fluorescence emission spectra recorded from isolated CP47 complexes of PSII ^32^. This shows that the presence and activity of LHCII significantly enhance photophysical processes that are already potentially present in PSII. On the one hand, recovering a certain fraction of the excitation energy already dissipated as heat shall increase the overall energy efficiency of PSII and therefore of photosynthesis. On the other hand, operation of this mechanism extends the action spectrum of PSII, towards the far-red spectral region, beyond the limit defined by the energy of the PSII reaction centre. A steep decrease in light absorption at wavelengths greater than 680 nm is associated with a very sharp decrease in the photosynthetic activity, referred to as the “red drop” ^10^. Importantly, a significant photosynthetic activity of light from the far-red spectral region has been demonstrated by direct measurements of photosynthetic oxygen evolution ^33, 34^ and by means of EPR spectroscopy ^35^. This effect has been observed with a single wavelength excitation ^34, 35^ and with isolated PSII particles ^35^ and therefore does not correspond directly to the classical Emerson effect ^36^. The photo-physical mechanism, reported in the present work, of thermally-driven up-conversion of the low-energy excitations to the energy level sufficiently high to drive the photochemical activity of PSII provides a direct explanation for the photosynthetic activity of light beyond the “red drop”, long-wavelength threshold. In our opinion, the mechanism of the thermally-driven up-conversion proved here to operate both in a model system and *in vivo*, is more realistic than involvement of unidentified “long-wavelength chlorophylls” ^34^ or hypothetical excited X* states ^35^. Importantly, the uphill activation energies for the far-red light-driven oxygen evolution in sunflower, determined based on the temperature dependencies, have been found to correspond directly to the energy gaps between the level attributed to PSII (680 nm) and the energies of light representing excitations in the long-wavelength spectral region ^34^. Interestingly, the quantum yield levels of PSII activity corresponding to 680 nm and the far-red region differ only by the factor of ~6, despite pronounced differences in light absorption in the corresponding spectral regions ^34^. A very close factor (6.3, see above) has been determined in the present work for the chlorophyll fluorescence quantum yield in LHCII, in the direct, downhill fluorescence excitation and the process of the uphill fluorescence excitation, mediated by the processes of thermally-driven up-conversion. The fact that two energetically coupled energy states reported in the present work, E680 and E700, are precisely tuned to the energies of the reaction centres of both photosystems, namely P680 and P700, provides favourable conditions for LHCII to act as a universal antenna complex. From the standpoint of energy supply to PSI, it can be assumed that there is no particular need to transfer low-energy excitations to this photosystem from LHCII. PSI of plants contains light absorption forms practically extending to 715 nm and emitting fluorescence in the 720-730 nm spectral region ^7^. Moreover, LHCII energetically coupled to PSI remains most probably in the trimeric form ^7^. Interestingly, the thermally-driven up-conversion has been recently shown to operate efficiently in isolated PSI ^37^. This can lead to a more general conclusion that uphill energy transfer is a common mechanism in the photosynthetic apparatus and potentially important for the process of photosynthesis. Natural photosynthesis constantly inspires technological activity focused on the construction of biomimetic solar cells based on isolated elements of the photosynthetic apparatus ^38^. The results of the present study show that engineering of solar cells containing not only photosynthetic reaction centers but additionally pigment-protein antenna complexes, in the form of supramolecular structures, can improve an overall energy yield of light energy conversion owing to the process of recycling of a fraction of excitation energy dissipated as heat. The results reported in the present study show that such a mechanism, shaped over millions of years of the biological evolution, is responsible for high activity of PSII in plants. Detailed excitation transfer pathways, involved in this mechanism, have been postulated in one selected system based on supramolecular structures of the major antenna complex LHCII. Such a description, based on selected structures of the photosynthetic apparatus, is naturally limited, but indicates the importance of the problem and sets directions for potential future research interests.

## Supporting information

Supporting Information (revised)

## ASSOCIATED CONTENT

### Supporting Information

The Supporting Information is available free of charge at https://

Experimental details, Excitation wavelength dependency of up-conversion-induced chlorophyll fluorescence in LHCII, Chlorophyll *a* fluorescence decay kinetics in LHCII, Fluorescence analysis of long-wavelength LHCII spectral forms, Table with fluorescence lifetime analysis of LHCII aggregated in the water phase (PDF).

## Author Contributions

^‖^M.Z. and R.L. contributed equally to this work.

## Notes

The authors declare no competing financial interest.

## ACKNOWLEDGEMENTS

The authors are grateful to prof. Pierre Joliot for stimulating discussion. We dedicate this work to Professor Joliot on the occasion of awarding him the title of Doctor Honoris Causa of the Maria Curie-Skłodowska University in Lublin. National Science Center of Poland is acknowledged for financial support within the project 2016/22/A/NZ1/00188. The research was carried out with the equipment purchased thanks to the financial support of the European Regional Development Fund in the framework of the Development of Eastern Poland Operational Program.

## REFERENCES

1. Hohmann-Marriott, M. F.; Blankenship, R. E., Evolution of Photosynthesis. Annu. Rev. Plant Biol. 2011, 62, 515–548.

2. Dau, H.; Zaharieva, I., Principles, Efficiency, and Blueprint Character of Solar-Energy Conversion in Photosynthetic Water Oxidation. Accounts Chem. Res. 2009, 42 (12), 1861–1870.

3. Joliot, P.; Joliot, A., A Polarographic Method for Detection of Oxygen Production and Reduction of Hill Reagent by Isolated Chloroplasts. Biochim. Biophys. Acta 1968, 153 (3), 625–634.

4. Blankenship, R. E., Origin and Early Evolution of Photosynthesis. Photosynth. Res. 1992, 33 (2), 91–111.

5. Rappaport, F.; Guergova-Kuras, M.; Nixon, P. J.; Diner, B. A.; Lavergne, J., Kinetics and Pathways of Charge Recombination in Photosystem II. Biochemistry 2002, 41 (26), 8518–8527.

6. Williamson, A.; Conlan, B.; Hillier, W.; Wydrzynski, T., The Evolution of Photosystem II: Insights Into the Past and Future. Photosynth. Res. 2011, 107 (1), 71–86.

7. Galka, P.; Santabarbara, S.; Thi, T. H. K.; Degand, H.; Morsomme, P.; Jennings, R. C.; Boekema, E. J.; Caffarri, S., Functional Analyses of the Plant Photosystem I-Light-Harvesting Complex II Supercomplex Reveal That Light-Harvesting Complex II Loosely Bound to Photosystem II Is a Very Efficient Antenna for Photosystem I in State II. Plant Cell 2012, 24 (7), 2963–2978.

8. Lamb, J. J.; Rokke, G.; Hohmann-Marriott, M. F., Chlorophyll fluorescence emission spectroscopy of oxygenic organisms at 77 K. Photosynthetica 2018, 56 (1), 105–124.

9. Jennings, R. C.; Belgio, E.; Zucchelli, G., Photosystem I, when excited in the chlorophyll Q(y) absorption band, feeds on negative entropy. Biophys Chem 2018, 233, 36–46.

10. Hall, D. O.; Rao, K. K.; Institute of Biology., Photosynthesis. 6th ed.; Cambridge University Press: Cambridge, UK; New York, 1999; p xiv, 214 p.

11. Liu, Z.; Yan, H.; Wang, K.; Kuang, T.; Zhang, J.; Gui, L.; An, X.; Chang, W., Crystal structure of spinach major light-harvesting complex at 2.72 A resolution. Nature 2004, 428 (6980), 287–92.

12. Standfuss, J.; Terwisscha van Scheltinga, A. C.; Lamborghini, M.; Kuhlbrandt, W., Mechanisms of photoprotection and nonphotochemical quenching in pea light-harvesting complex at 2.5 A resolution. EMBO J. 2005, 24 (5), 919–928.

13. Gruszecki, W. I.; Grudzinski, W.; Gospodarek, M.; Patyra, M.; Maksymiec, W., Xanthophyll-induced aggregation of LHCII as a switch between light-harvesting and energy dissipation systems. Biochim. Biophys. Acta 2006, 1757 (11), 1504–1511.

14. Johnson, M. P.; Goral, T. K.; Duffy, C. D. P.; Brain, A. P. R.; Mullineaux, C. W.; Ruban, A. V., Photoprotective Energy Dissipation Involves the Reorganization of Photosystem II Light-Harvesting Complexes in the Grana Membranes of Spinach Chloroplasts. Plant Cell 2011, 23 (4), 1468–1479.

15. Kirchhoff, H., Molecular crowding and order in photosynthetic membranes. Trends Plant Sci. 2008, 13 (5), 201–207.

16. Natali, A.; Gruber, J. M.; Dietzel, L.; Stuart, M. C. A.; van Grondelle, R.; Croce, R., Light-harvesting Complexes (LHCs) Cluster Spontaneously in Membrane Environment Leading to Shortening of Their Excited State Lifetimes. J. Biol. Chem. 2016, 291 (32), 16730–16739.

17. Akhtar, P.; Gorfol, F.; Garab, G.; Lambrev, P. H., Dependence of Chlorophyll Fluorescence Quenching on the Lipid-to-Protein Ratio in Reconstituted Light-Harvesting Complex II Membranes Containing Lipid Labels. Chem. Phys. 2019, 522, 242–248.

18. Horton, P.; Ruban, A. V.; Rees, D.; Pascal, A. A.; Noctor, G.; Young, A. J., Control of the light-harvesting function of chloroplast membranes by aggregation of the LHCII chlorophyll-protein complex. FEBS Lett. 1991, 292, 1–4.

19. Janik, E.; Bednarska, J.; Zubik, M.; Luchowski, R.; Mazur, R.; Sowinski, K.; Grudzinski, W.; Garstka, M.; Gruszecki, W. I., A chloroplast "wake up" mechanism: Illumination with weak light activates the photosynthetic antenna function in dark-adapted plants. J. Plant Physiol. 2017, 210, 1–8.

20. Janik, E.; Bednarska, J.; Zubik, M.; Puzio, M.; Luchowski, R.; Grudzinski, W.; Mazur, R.; Garstka, M.; Maksymiec, W.; Kulik, A.; Dietler, G.; Gruszecki, W. I., Molecular architecture of plant thylakoids under physiological and light stress conditions: A study of lipid-light-harvesting complex II model membranes. Plant Cell 2013, 25, 2155–2170.

21. Zhou, J.; Sekatskii, S.; Welc, R.; Dietler, G.; Gruszecki, W. I., The role of xanthophylls in the supramolecular organization of the photosynthetic complex LHCII in lipid membranes studied by high-resolution imaging and nanospectroscopy. Biochim Biophys Acta Bioenerg 2020, 1861 (2), 148117.

22. Chmeliov, J.; Gelzinis, A.; Songaila, E.; Augulis, R.; Duffy, C. D. P.; Ruban, A. V.; Valkunas, L., The nature of self-regulation in photosynthetic light-harvesting antenna. Nat Plants 2016, 2 (5), 16045.

23. Ruban, A. V.; Dekker, J. P.; Horton, P.; Van Grondelle, R., Temperature-Dependence of Chlorophyll Fluorescence from the Light-Harvesting Complex Ii of Higher-Plants. Photochem. Photobiol. 1995, 61 (2), 216–221.

24. Zubik, M.; Luchowski, R.; Puzio, M.; Janik, E.; Bednarska, J.; Grudzinski, W.; Gruszecki, W. I., The negative feedback molecular mechanism which regulates excitation level in the plant photosynthetic complex LHCII: Towards identification of the energy dissipative state. Biochim. Biophys. Acta 2013, 1827 (3), 355–364.

25. Belgio, E.; Johnson, M. P.; Juric, S.; Ruban, A. V., Higher Plant Photosystem II Light-Harvesting Antenna, Not the Reaction Center, Determines the Excited-State Lifetime-Both the Maximum and the Nonphotochemically Quenched. Biophys. J. 2012, 102 (12), 2761–2771.

26. Toth, T. N.; Rai, N.; Solymosi, K.; Zsiros, O.; Schroder, W. P.; Garab, G.; van Amerongen, H.; Horton, P.; Kovacs, L., Fingerprinting the macro-organisation of pigment-protein complexes in plant thylakoid membranes in vivo by circular-dichroism spectroscopy. Bba-Bioenergetics 2016, 1857 (9), 1479–1489.

27. Yamamoto, Y.; Hori, H.; Kai, S.; Ishikawa, T.; Ohnishi, A.; Tsumura, N.; Morita, N., Quality control of Photosystem II: reversible and irreversible protein aggregation decides the fate of Photosystem II under excessive illumination. Front Plant Sci 2013, 4.

28. Kruger, T. P. J.; Ilioaia, C.; Johnson, M. P.; Ruban, A. V.; van Grondelle, R., Disentangling the low-energy states of the major light-harvesting complex of plants and their role in photoprotection. Bba-Bioenergetics 2014, 1837 (7), 1027–1038.

29. Van Grondelle, R., Excitation-Energy Transfer, Trapping and Annihilation in Photosynthetic Systems. Biochim. Biophys. Acta 1985, 811 (2), 147–195.

30. Akhtar, P.; Do, T. N.; Nowakowski, P. J.; Huerta-Viga, A.; Khyasudeen, M. F.; Lambrev, P. H.; Tan, H. S., Temperature Dependence of the Energy Transfer in LHCII Studied by Two-Dimensional Electronic Spectroscopy. J Phys Chem B 2019, 123 (31), 6765–6775.

31. Gruszecki, W. I.; Luchowski, R.; Zubik, M.; Grudzinski, W., Photothermal Microscopy: Imaging of Energy Dissipation From Photosynthetic Complexes. Anal. Chem. 2015, 87 (19), 9572–9575.

32. Andrizhiyevskaya, E. G.; Chojnicka, A.; Bautista, J. A.; Diner, B. A.; van Grondelle, R.; Dekker, J. P., Origin of the F685 and F695 Fluorescence in Photosystem II. Photosynth. Res. 2005, 84 (1-3), 173–180.

33. Pettai, H.; Oja, V.; Freiberg, A.; Laisk, A., The Long-Wavelength Limit of Plant Photosynthesis. FEBS Lett. 2005, 579 (18), 4017–4019.

34. Pettai, H.; Oja, V.; Freiberg, A.; Laisk, A., Photosynthetic Activity of Far-Red Light in Green Plants. Biochim. Biophys. Acta 2005, 1708 (3), 311–321.

35. Thapper, A.; Mamedov, F.; Mokvist, F.; Hammarstrom, L.; Styring, S., Defining the Far-Red Limit of Photosystem II in Spinach. Plant Cell 2009, 21 (8), 2391–2401.

36. Emerson, R., Dependence of Yield of Photosynthesis in Long-Wave Red on Wavelength and Intensity of Supplementary Light. Science 1957, 125 (3251), 746–746.

37. Giera, W.; Szewczyk, S.; McConnell, M. D.; Redding, K. E.; van Grondelle, R.; Gibasiewicz, K., Uphill energy transfer in photosystem I from Chlamydomonas reinhardtii. Time-resolved fluorescence measurements at 77 K. Photosynth. Res. 2018, 137 (2), 321–335.

38. Ravi, S. K.; Tan, S. C., Progress and Perspectives in Exploiting Photosynthetic Biomolecules for Solar Energy Harnessing. Energ. Environ. Sci. 2015, 8 (9), 2551–2573.

