## Supporting Information (revised) for "Recycling of energy dissipated as heat accounts for high activity of Photosystem II"

### Experimental Section

*Plant material and treatment*

*Arabidopsis thaliana* (L.) wild-type Columbia 406 (Col-0) seeds were obtained from the Nottingham Arabidopsis Stock Centre (NASC). Plants used in the experiments were grown in a growth chamber under controlled environmental conditions with air humidity at 60 %, with day and night temperatures of 22°C and 18°C, respectively. Plants were grown in a mixture of soil and sand of optimum soil water availability controlled by daily weighing and watering. The average light intensity at the level of rosette was maintained at 150 μmol photons $m^{-2}$ $s^{-1}$ during an 8-hr light cycle. Measurements were carried out on mature leaves of 8-week-old plants.

In order to obtain intact cells with chloroplasts, the bottom epidermis from *Arabidopsis thaliana* leaves was peeled off using Scotch tape (3M). The sample was placed onto non-fluorescent glass slides (Menzel-Glaser), soaked in a buffer (20 mM Tricine, 0.4 M Sorbitol, pH 7.8) and subjected to measurements.

*Preparation of lipid-LHCII membranes and aggregated structures of LHCII*

LHCII was isolated according to [1] with slight modifications [2]. The purity of preparations was controlled using HPLC and electrophoretic methods [2]. Lipid-LHCII membranes were prepared according to [3]. MGDG and DGDG (Avanti Polar Lipids Inc., USA) were mixed in chloroform:methanol (2:1, v:v) solution in a molar ratio of 2:1. Next, the mixture was dried under a stream of nitrogen to obtain a thin film in a glass tube. Obtained samples were placed in a vacuum ($10^{-5}$ bar) for 40 min in order to remove traces of organic solvents. For incorporation of LHCII into lipid membranes, LHCII complexes suspended in tricine buffer (20 mM Tricine, 10 mM KCl, pH 7.6) containing 0.1% n-dodecyl-β-D-maltoside (DM) were transferred to glass tubes containing the deposited lipid film and subjected to mild sonication using an ultrasonic bath for 30 min. The molar ratio of LHCII:lipids was 1:200. Detergent (DM) was removed from the suspension by incubation with Bio-beads adsorbent (Bio-Rad Laboratories, USA) at 4℃ for 14 h. The pellet obtained by centrifugation for 5 min at 14000 x g consisted of the lipid-LHCII membranes.

Sample containing aggregated structures of LHCII in the model system without lipids was prepared following the same protocol except that the protein suspension in 0.1 % DM has not been mixed with lipids but subjected directly to incubation with Bio-beads detergent adsorbent.

*Fluorescence spectroscopy*

Fluorescence emission spectra of leaves and LHCII samples were recorded with an FS5 spectrofluorometer (Edinburgh Instruments, UK) with excitation and emission bandwidths set to 3 nm. In order to avoid spectral distortion caused by the Kautsky effect, each leaf was preilluminated with a white light LED lamp with the controlled photon flux density (150 μmol photons $m^{-2}$ $s^{-1}$, 2 min) prior to the recording of fluorescence spectra.

Time-resolved fluorescence intensity decays were measured using FluoTime 300 spectrometer (PicoQuant, Germany). Excitation at 635 nm with 20 MHz frequency of pulses was from solid-state laser LDH-P-635 with a pulse width 68 ps, 214 μW. Excitation at 705 nm with 4 MHz repetition rate, 1.03 mW was accomplished by a single pulse selector (APE, GmbH) with a tunable Ti-sapphire laser Chameleon Ultra (Coherent, Inc.). The emission at 680 nm was filtered by 680/13 bandpass filter (Semrock, Inc.) and 665 glass long-wavelength pass filter (Edmund Optics, Inc.). Detection was performed with a micro-channel plate and time-correlated single-photon counting system PicoHarp 300. Fluorescence lifetime decays were fitted using FluoFit Pro software (PicoQuant, Germany). A FluoTime 300 spectrometer was also used to record fluorescence emission spectra of samples excited at 635 nm and 705 nm, with the application of the lasers specified above. The fluorescence quantum yields, excited at those wavelengths, were compared based on the integration of the emission spectra and taking into consideration the laser powers and the values of light absorption by the sample at selected wavelengths (calculated as 1-transmission). Steady-state and time-resolved fluorescence measurements at different temperatures were conducted with the application of an Optistat DN2 Cryostat (Oxford Instruments, UK). The LHCII samples to be measured in a frozen state were diluted in buffer-glycerol (1:2, v:v). An important issue in the recording and analysis of chlorophyll fluorescence spectra *in vivo* is a proper correction for possible light reabsorption [4]. In order to minimize spectral distortions due to fluorescence reabsorption in leaves, Chl *a* fluorescence emission spectra *in vivo* were recorded from the epidermis separated from a leaf and placed in a cryostat. In order to correct such emission spectra for the effect of fluorescence reabsorption, each emission spectrum was divided by the transmission spectrum recorded from the same sample, under identical experimental conditions. Such an approach certainly does not lead to underestimation the fluorescence reabsorption effect in the spectral region characteristic of light emission by Chl *a* molecules associated with PSII.

REFERENCES

1. Krupa, Z.; Huner, N. P. A.; Williams, J. P.; Maissan, E.; James, D. R., Development at cold-hardening temperatures. The structure and composition of purified rye light harvesting complex II. *Plant Physiol.* 1987, *84*, 19-24.

2. Gruszecki, W. I.; Gospodarek, M.; Grudzinski, W.; Mazur, R.; Gieczewska, K.; Garstka, M., Light-induced Change of Configuration of the LHCII-Bound Xanthophyll (Tentatively Assigned to Violaxanthin): A Resonance Raman Study. *J. Phys. Chem. B* 2009, *113* (8), 2506-2512.

3. Janik, E.; Bednarska, J.; Zubik, M.; Puzio, M.; Luchowski, R.; Grudzinski, W.; Mazur, R.; Garstka, M.; Maksymiec, W.; Kulik, A.; Dietler, G.; Gruszecki, W. I., Molecular

architecture of plant thylakoids under physiological and light stress conditions: A study of lipid-light-harvesting complex II model membranes. *Plant Cell* 2013, *25*, 2155-2170.

4. Weis, E., Chlorophyll Fluorescence at 77-K in Intact Leaves - Characterization of a Technique to Eliminate Artifacts Related to Self-Absorption. *Photosynth Res* 1985, *6* (1), 73-86.

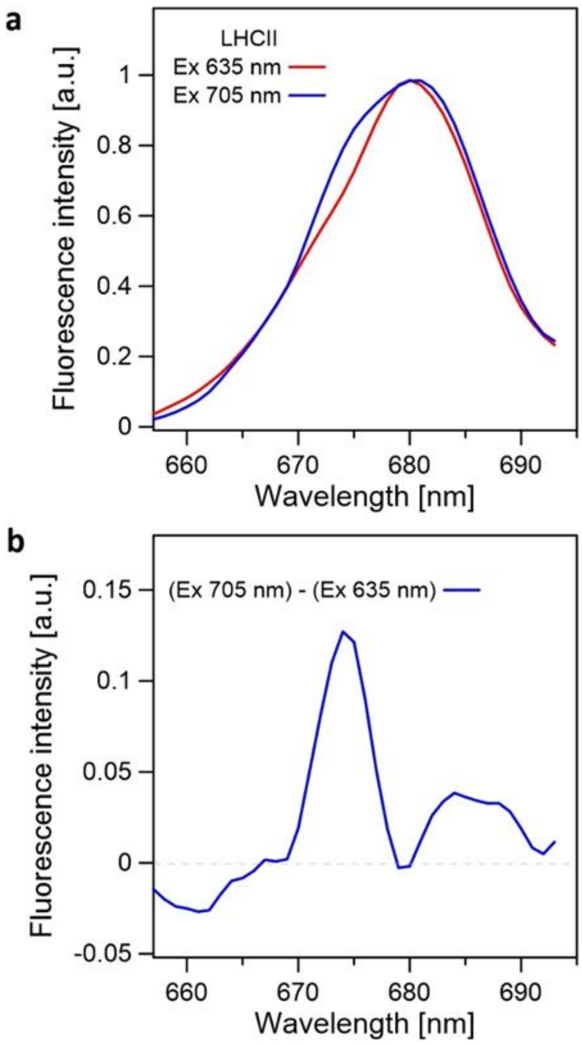


**Figure S1.** Fluorescence emission spectra of chlorophyll *a* in LHCII. Panel a presents the spectra displayed in Fig. 1c of the paper, recorded with excitation wavelength at 635 nm and 705 nm (marked). Panel b presents the difference spectra calculated by subtraction of the spectra presented in panel a: recorded with excitation at 705 nm minus the spectrum recorded with excitation at 635 nm. The difference spectrum shows a distinct fluorescence emission band centering at 673 nm and an additional band at 685 nm. Such an effect can be interpreted in terms of heterogeneity of spectral forms of LHCII, spontaneously formed in the chloroplast lipid system, some of which demonstrate particularly efficient thermally-driven up-conversion to the electronic states characterized by the fluorescence bands centering at 673 nm and 685 nm. In contrast, such an effect attributed to a heterogeneity of spectral forms has not been observed in the aggregated structures of LHCII formed in the water phase (see Fig. S6).

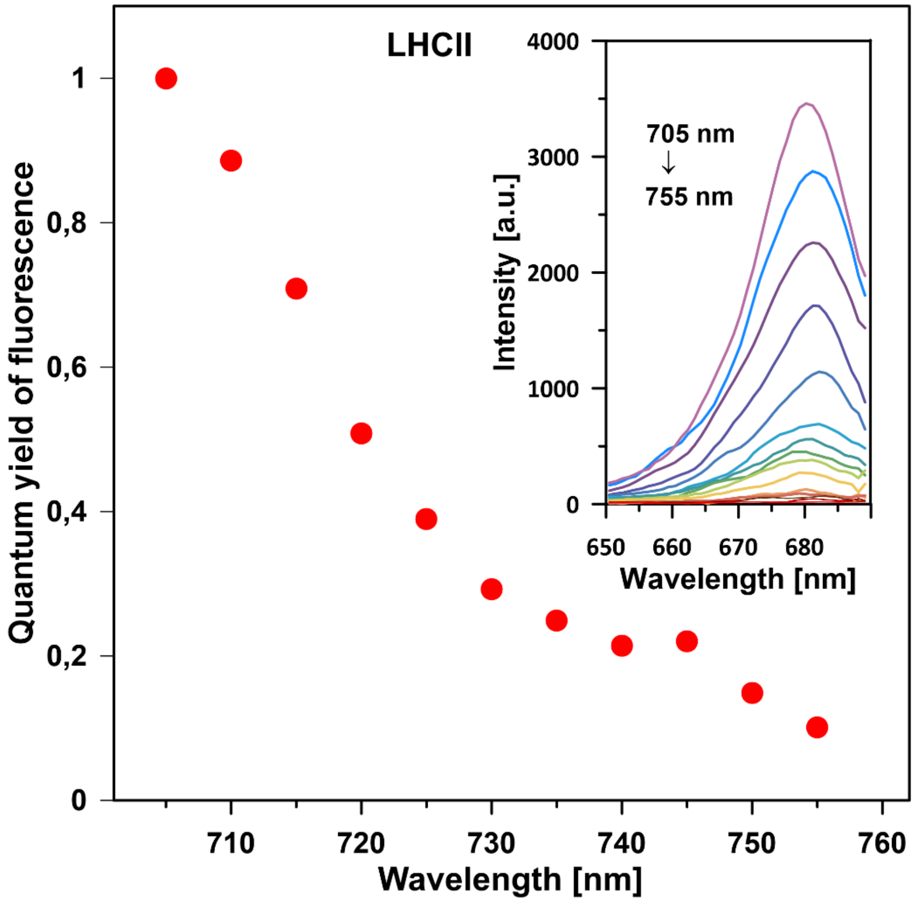


**Figure S2.** Excitation wavelength dependency of up-conversion-induced chlorophyll fluorescence in LHCII. Chlorophyll *a* fluorescence in LHCII-lipid membranes was excited at different wavelengths and detected in the spectral window presented in the inset. The spectra were recorded at 297 K. The experimental points in the main graph represent the relative fluorescence quantum yield values (normalized at the maximum) calculated based on the integration of the emission spectra, 1 minus transmission (1-T) spectrum of the sample and a number of photons emitted by the tunable laser at each excitation wavelength.

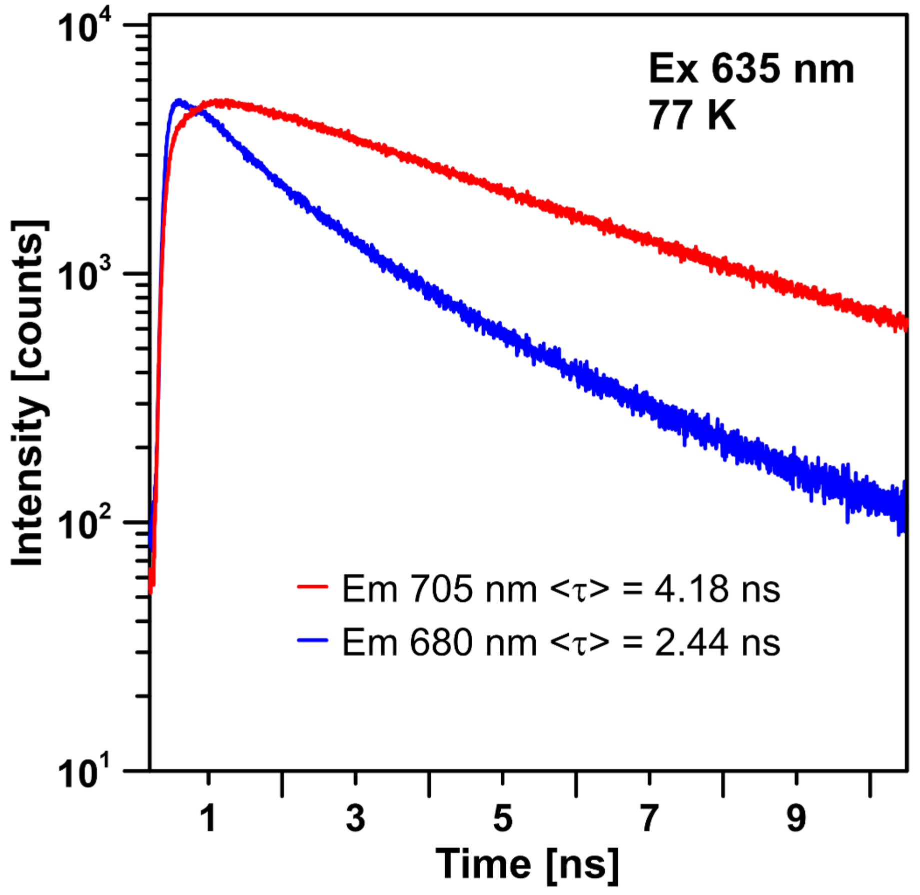


**Figure S3.** Chlorophyll *a* fluorescence decay kinetics in LHCII. Fluorescence decay traces recorded from LHCII-lipid membranes excited at 635 nm and observed either at 705 nm or at 680 nm (indicated). Sample temperature 77 K. The average fluorescence lifetimes determined based on the decays presented in this figure are reported on the diagram presented in Figure 3 of the paper.

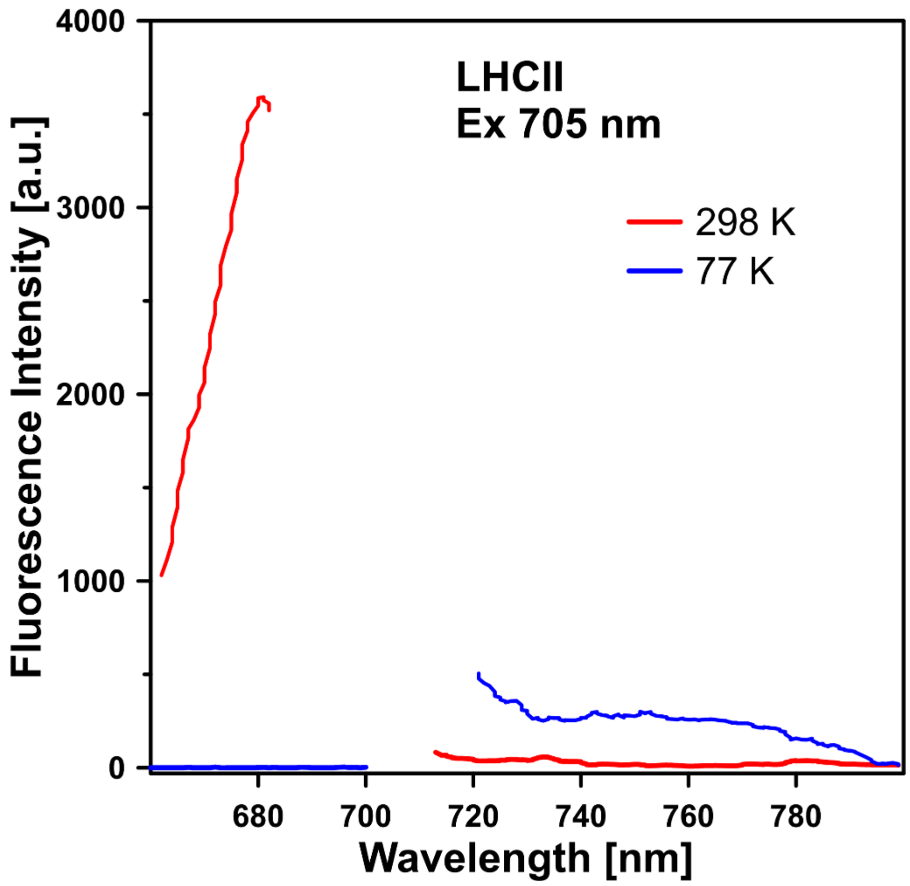


**Figure S4.** Fluorescence emission spectra of chlorophyll *a* in LHCII. The spectra were recorded from LHCII-lipid membranes either at 77 K or at 298 K. The sample containing aggregated LHCII was excited at 705 nm and the emission spectra were recorded in the lower energy spectral region as well as in the higher energy spectral region, both at 298 K and 77 K. Note that at 298 K the photons absorbed are almost exclusively emitted in the short-wavelength region, in the consequence of the thermally-driven up-conversion, in contrast to the measurement conducted at 77 K. Under such conditions, fluorescence is exclusively emitted in the longer-wavelength spectral window.

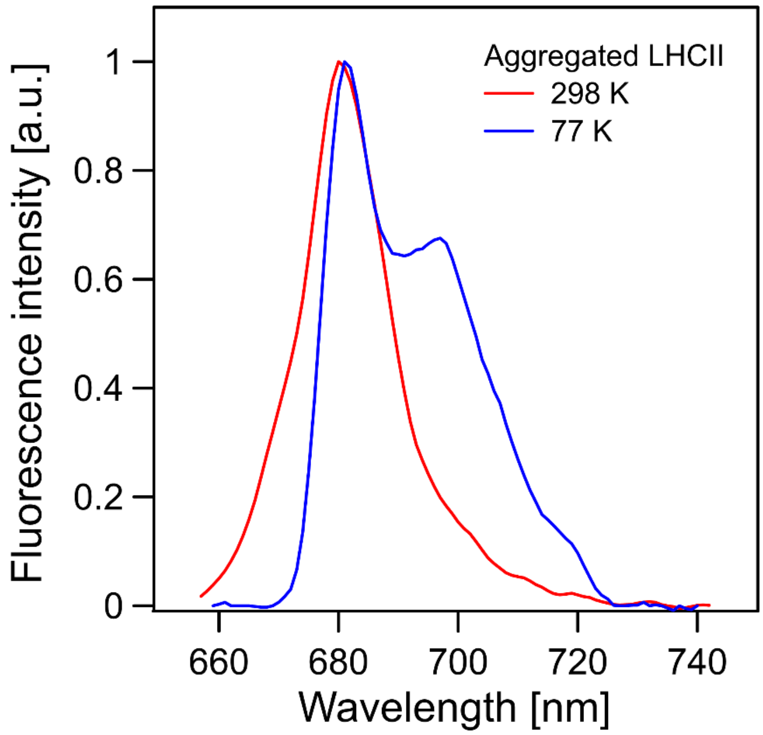


**Figure S5**. Fluorescence emission spectra of chlorophyll *a* in aggregated LHCII. Aggregation of LHCII was achieved by elimination of a detergent from the protein suspension in a water medium. Fluorescence emission spectra recorded at 77 K and 298 K (indicated) with excitation at 635 nm. High intensity of the spectral component in the region of 700 nm, relative to the principle spectral component in the region of 680 nm, is a manifestation of the presence of aggregated structures of LHCII.

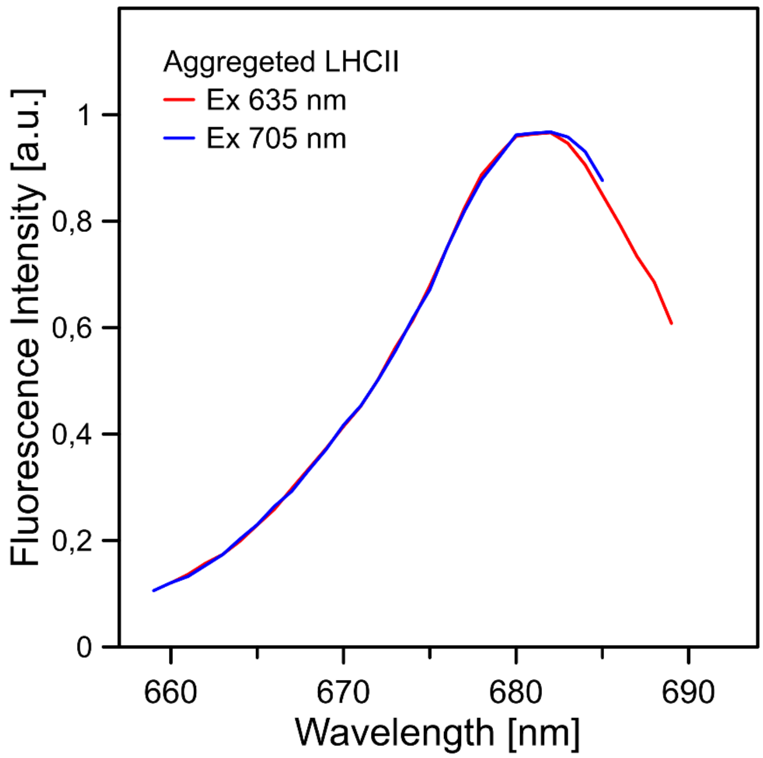


**Figure S6**. Fluorescence emission spectra of chlorophyll *a* in aggregated LHCII. Aggregation of LHCII was achieved by elimination of a detergent from the protein suspension in a water medium. Fluorescence emission spectra recorded at 298 K with excitation at 635 nm or at 705 nm (indicated).

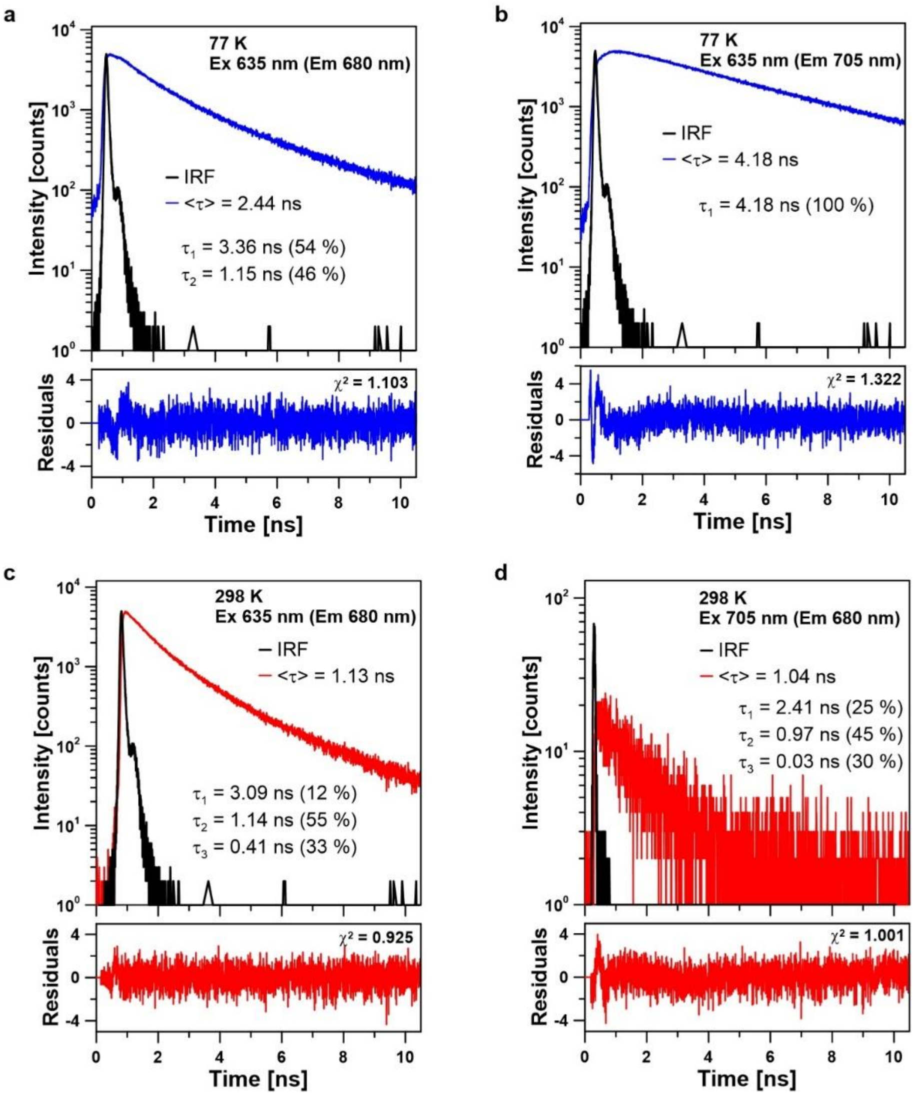


**Figure S7.** Chlorophyll *a* fluorescence decay kinetics in LHCII. Fluorescence decay traces recorded from LHCII-lipid membranes were recorded at 77 K (upper panels) or 298 K (lower panels). Excitation and emission wavelengths are reported in each panel. The decay kinetics were analyzed within the framework of multiexponential functions, with a component number between 1 and 3, dependently on a criterion based on the wellness of fit (presented beneath each panel). Fluorescence lifetime components determined are presented in each panel along with relative amplitudes. The average fluorescence lifetimes determined based on the decays presented in this figure are reported on the diagram presented in Figure 3 of the paper.

**Table S1**

Fluorescence lifetime parameters of Chl *a* in LHCII aggregated in the water environment.

| Temperature [K] | Excitation wavelength [nm] | Emission wavelength [nm] | Average fluorescence lifetime < τ > [ns] | Lifetime component $\tau_1$ [ns] | Lifetime component $\tau_2$ [ns] |
|---|---|---|---|---|---|
| **298** | **635** | **680** | 0.16 | 0.23 (31 %) | 0.13 (69 %) |
| | | **700** | 0.18 | 0.15 (88 %) | 0.35 (12 %) |
| | **705** | **680** | 0.14 | 0.19 (66 %) | 0.04 (34 %) |
| **77** | **635** | **680** | 2.99 | 1.17 (20%) | 3.45 (80 %) |
| | | **700** | 3.14 | 1.16 (49%) | 3.74 (51 %) |
| | **705** | **680** | n.d. | n.d. | n.d. |
